# Demonstration of electron diffraction from membrane protein crystals grown in a lipidic mesophase after lamella preparation by focused ion beam milling at cryogenic temperatures

**DOI:** 10.1101/2020.07.03.186049

**Authors:** Vitaly Polovinkin, Krishna Khakurel, Michal Babiak, Borislav Angelov, Bohdan Schneider, Jan Dohnalek, Jakob Andreasson, Janos Hajdu

## Abstract

Electron crystallography of sub-micron sized 3D protein crystals has emerged recently as a valuable field of structural biology. *In meso* crystallization methods, utilizing lipidic mesophases, particularly lipidic cubic phases (LCPs), can produce high-quality 3D crystals of membrane proteins (MPs). A major step towards realising 3D electron crystallography of MP crystals, grown *in meso*, is to demonstrate electron diffraction from such crystals. The first task is to remove the viscous and sticky lipidic matrix, surrounding the crystals without damaging the crystals. Additionally, the crystals have to be thin enough to let electrons traverse them without significant multiple scattering. In the present work, we experimentally verified the concept that focused ion beam milling at cryogenic temperatures (cryo-FIB) can be used to remove excess host lipidic mesophase matrix, and then thin the crystals to a thickness suitable for electron diffraction. In this study, bacteriorhodopsin (BR) crystals grown in a lipidic mesophase of monoolein were used as a model system. LCP from a part of a 50-μm thick crystal, which was flash-frozen in liquid nitrogen, was milled away with a gallium FIB under cryogenic conditions, and a part of the crystal itself was thinned into a ∼210-nm thick lamella with the ion beam. The frozen sample was then transferred into an electron cryo-microscope (cryo-EM), and a nanovolume of ∼1400×1400×210 nm^3^ of the BR lamella was exposed to 200-kV electrons at a fluence of ∼0.06 e^−^/Å^2^. The resulting electron diffraction peaks were detected beyond 2.7-Å resolution (with mean signal-to-noise ratio <I/σ(I)> of >7) by a CMOS-based Ceta 16M camera. The results demonstrate, that cryo-FIB milling produces high quality lamellae from crystals grown in lipidic mesophases, and pave the way for 3D electron crystallography on crystals grown or embedded in highly viscous media.

**Synopsis:** Electron diffraction experiments on crystals of membrane proteins grown in lipidic mesophases have not been possible due to a thick layer of viscous crystallisation medium around the crystals. Here we show that focused ion beam milling at cryogenic temperatures (cryo-FIB milling) can remove the viscous layer, and demonstrate high-quality electron diffraction on a FIB-milled lamella of a bacteriorhodopsin 3D crystal.

## 1. Introduction

Electron crystallography of three-dimensional (3D) macromolecular crystals is a promising technique for solving structures at high resolution (Nannenga & Gonen, 2019). Profiting from strong interactions of electrons with matter (Henderson, 1995) and utilizing a standard cryo-electron microscope (cryo-EM) (Shi *et al*., 2013, 2016), electron diffraction provides an effective means of solving structures of proteins and peptides from very thin crystals, which would either be difficult to study by synchrotron-based methods or can only be investigated with X-ray free-electron lasers (XFELs) (Neutze *et al*., 2000; Chapman *et al*., 2011; Wolff *et al*., 2020). Moreover, atomic scattering factors for electrons, unlike those for X-rays, are strongly influenced by the charge state of the scattering atoms. This makes electron crystallography particularly suitable for mapping charged states of residues and cofactors of biomolecules and, in general, for investigating charge transfer and ion transport processes in biochemical reactions (Kimura *et al*., 1997; Mitsuoka *et al*., 1999; Subramaniam & Henderson, 2000; Gadsby, 2009; Yonekura *et al*., 2015; Sjulstok *et al*., 2015).

Since 2013, when the first protein structure was determined by the microcrystal electron diffraction (MicroED) technique (Shi *et al*., 2013), 3D macromolecular crystallography with electrons has undergone significant developments in data collection and data analysis (Nannenga *et al*., 2014; van Genderen *et al*., 2016; Clabbers *et al*., 2017; Gruene *et al*., 2018; Hattne *et al*., 2019; Bücker *et al*., 2020), and a number of new protein and peptide structures have been solved (Rodriguez *et al*., 2015; de la Cruz *et al*., 2017; Xu *et al*., 2019). For a typical ED data collection experiment, crystals in solution are pipetted onto a transmission electron microscopy (TEM) grid, blotted to remove excess solution, and vitrified in liquid ethane or nitrogen. Finally, the diffraction data are recorded in a movie mode as the crystals are continuously rotated under parallel illumination with a high-energy electron beam (typically, 200-kV) in a transmission cryo-EM (Shi *et al*., 2016). The crystal thickness along the incident beam should be of sub-micron dimensions to reduce multiple scattering and absorption (Zeldin & Brunger, 2013; Vulović *et al*., 2013; Yan *et al*., 2015; Subramanian *et al*., 2015).

Protein and peptide crystals below ∼0.5-µm thickness have successfully been used for structure determination (Rodriguez *et al*., 2015; Sawaya *et al*., 2016; Clabbers *et al*., 2017; Xu *et al*., 2019). Bigger crystals can be broken up to smaller fragments by mechanical force (e.g. vigorous pipetting, sonication, vortexing with beads and etc.) (de la Cruz *et al*., 2017) or can be thinned in a more reproducible and reliable way to a desired size by focused ion beam milling under cryogenic conditions (cryo-FIB) (Duyvesteyn *et al*., 2018; Zhou *et al*., 2019; Martynowycz *et al*., 2019*a*). The latter technique has been inherited from material science (Giannuzzi & Stevie, 1999) and cellular cryo-electron tomography (cryo-ET) (Marko *et al*., 2007; Plitzko & Baumeister, 2019). Cryo-FIB milling enables ∼10-nm precision in the production of crystal lamellae of hundreds of nanometers in thickness and thus provides an important means of optimizing the ED signal (Zhou *et al*., 2019). As cryo-FIB instrumentation becomes more accessible, this technique can become a standard tool, extending the capabilities of electron crystallography (Zuo & Spence, 1992; Stokes *et al*., 2013; de la Cruz *et al*., 2017; Nannenga & Gonen, 2019; Nannenga, 2020). We show here that cryo-FIB milling can overcome sample preparation challenges specific to membrane protein crystals grown in a viscous environment.

*In meso* crystallisation of membrane proteins (MP) utilizes lipidic mesophases (Ishchenko *et al*., 2017; Caffrey & Cherezov, 2009), including lipidic cubic phases (LCPs) (Landau & Rosenbusch, 1996), where crystals grow in an extremely viscous medium, with viscosities up to ∼50 Pa·s (Perry *et al*., 2009). The viscous crystallisation medium cannot be easily removed from such crystals on TEM grids (Shi *et al*., 2016; de la Cruz *et al*., 2017), and this poses problems for electron diffraction due to the low penetration depth of electrons, which may be stopped in the bulky lipidic mesophase. The penetration depth of X-rays is orders of magnitude higher than that of electrons, and so the LCP technique has been successful in X-ray studies (see e.g. (Pebay-Peyroula *et al*., 1997; Gordeliy *et al*., 2002; Cherezov *et al*., 2007; Caffrey, 2015)). In contrast, not a single MP structure has been reported by electron diffraction from crystals grown *in meso*. The layer of the viscous crystallisation medium can be several microns thick over the crystal, hindering the propagation of electrons. In principle, one can lower the viscosity of the LCP environment or dissolve it in a “chemical” way by adding detergents (Luecke *et al*., 1999), so-called spongifiers (Cherezov *et al*., 2006), oils (Niwa & Takeda, 2019) or by treating the LCP matrix with a lipase (Belrhali *et al*., 1999) and then use standard blotting techniques to remove the liquid. This chemical approach was successfully demonstrated on non-membrane proteins, e.g. proteinase K crystals embedded into a LCP matrix (Zhu *et al*., 2019). The chemical process is sample-dependent and requires screening of phase-dissolving conditions to leave the particular type of crystal intact.

We demonstrate here that FIB milling at cryogenic temperatures has the potential to become a universal alternative to chemical cleaning. Conceptually, MP crystals grown in a lipidic mesophase are deposited (or grown directly) on a TEM grid, then vitrified by plunge-freezing, and the excess of lipidic crystallization matrix is milled away by the cryo-FIB. The thickness of the crystal can be adjusted to the desired level, to enable penetration of the electron beam and observation of the diffraction signal. This procedure can be general, unlike chemical cleaning, and could become a universal step in sample preparation for ED. The technique can potentially be applied to membrane or water-soluble protein crystals grown/embedded in a lipidic mesophase or, in general, in a viscous crystallization medium (e.g. bicelles, vesicles (Ishchenko *et al*., 2017)). Here, we report the first observed ED diffraction signal from a 3D MP crystal grown in a lipidic cubic phase. The crystal was cleaned up from the crystallization matrix and milled down to a thin lamella (∼210-nm, in this case), using the cryo-FIB.

## 2. Material and methods

### 2.1. Protein crystallization

Expression, purification, solubilisation in β-octyl glucoside (Cube Biotech) and crystallization of BR (from *Halobacterium salinarum S9* cells) in the lipidic cubic phase of monoolein (Nu-Chek Prep) were carried out as described in (Gordeliy *et al*., 2003). Aliquots (200 nl) of a protein-mesophase mixture were spotted on a 96-well lipidic cubic phase glass sandwich plate (Marienfeld) and overlaid with 800 nl of precipitant solution by means of an NT8 crystallization robot (Formulatrix). Na/K-phosphate (pH 5.6) solutions at concentrations from 1 to 2.4 M (in concentration of phosphate) were used as precipitant. Hexagonally shaped plate-like BR crystals of space group P6_3_ were observed after 2–3 weeks at room temperature.

### 2.2. Sample preparation for ED experiments by cryo-FIB milling

The sample preparation workflow is schematically shown in Fig.1. The crystals were harvested directly from the LCP matrix (Fig. 2(a)) using a MiTeGen microloop, then positioned on a holey carbon TEM grid (Quantifoil R2/1, Cu, 200-mesh) pre-clipped into an Autogrid assembly (Thermo Fisher Scientific) and manually plunge-frozen in liquid nitrogen. The frozen sample was transferred into a dual-beam focused ion beam scanning electron microscope (FIB-SEM) (Versa 3D, Thermo Fisher Scientific), equipped with the Quorum PP3010T cryogenic sample preparation-transfer system. The specimen was kept at −192°C on the cryo-stage during the whole procedure. The lamella preparation protocol followed the approach published earlier (see (Schaffer *et al*., 2017; Martynowycz *et al*., 2019*b*; Zhou *et al*., 2019)). The crystal was coated with a conductive Pt–sputter (10 mA, 60 s) layer and a protective organometallic Pt-GIS (30°C, 80 s) layer. The crystal was milled by rastering over the area of interest (Fig. 2(d)), slowly removing portions of the sample above and below the selected area. Milling was performed in a stepwise way, using a 30-kV gallium ion (Ga^+^) beam and reducing the beam current as the crystal lamella was thinned down to a final thickness of ∼210 nm (Fig. 3). Currents of 1 nA, 300 pA, 100 pA, 30 pA and 10 pA were used (in this order) to generate a final lamella by decreasing the thickness of the sample to ∼5 μm, ∼2 μm, ∼1 μm, ∼400 nm and all the way to the final thickness of ∼210 nm, respectively. The lamella was produced at an angle of 22° with respect to the grid plane. The grid with the BR crystal lamella was transferred to a TEM microscope (Talos Arctica, Thermo Fisher Scientific) for the collection of the ED data.

**Figure 1.**
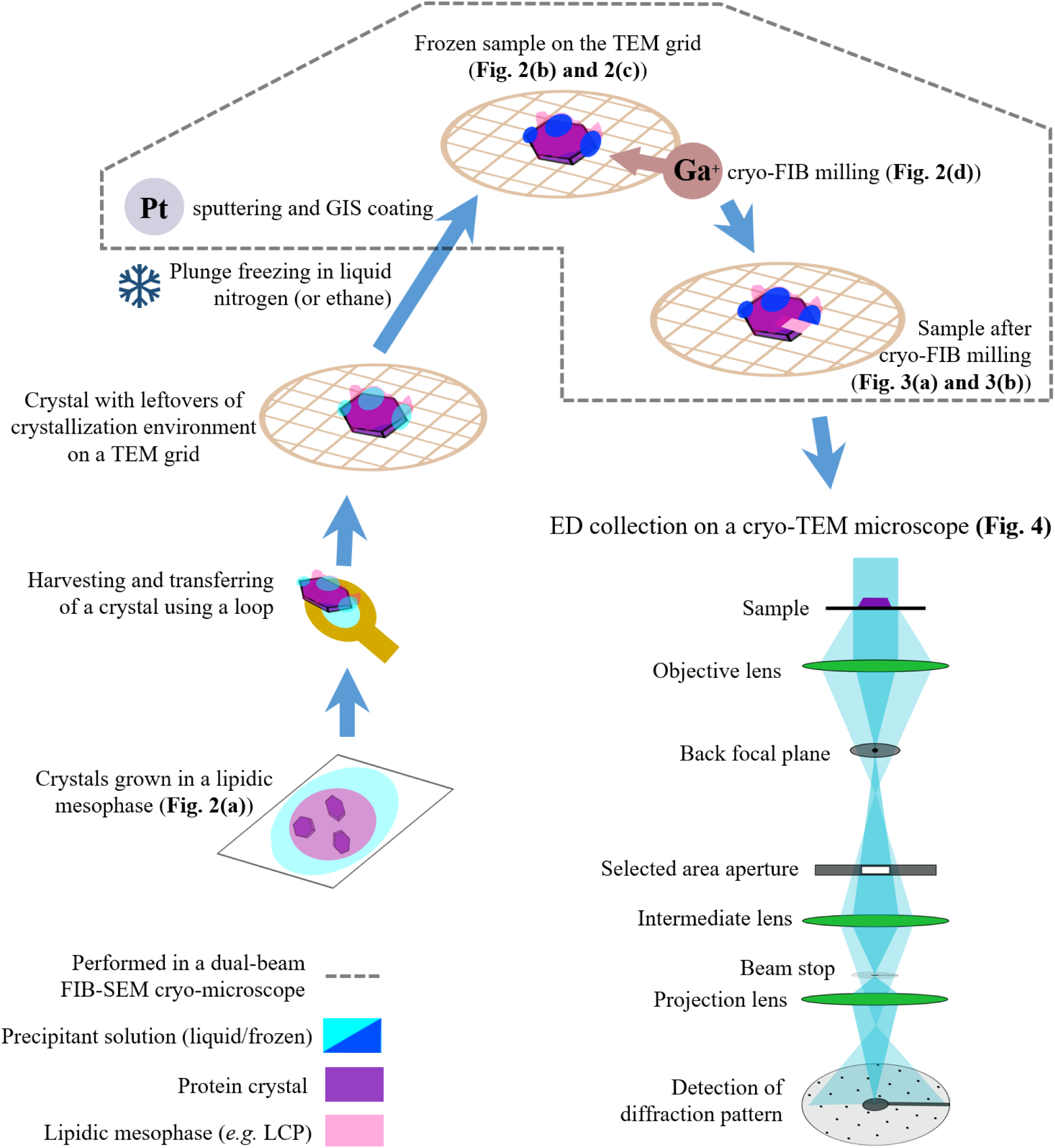
Schematic diagram of the experimental workflow. Schematic view of the electron diffraction setup is adapted from (Rodriguez & Gonen, 2016).

**Figure 2.**
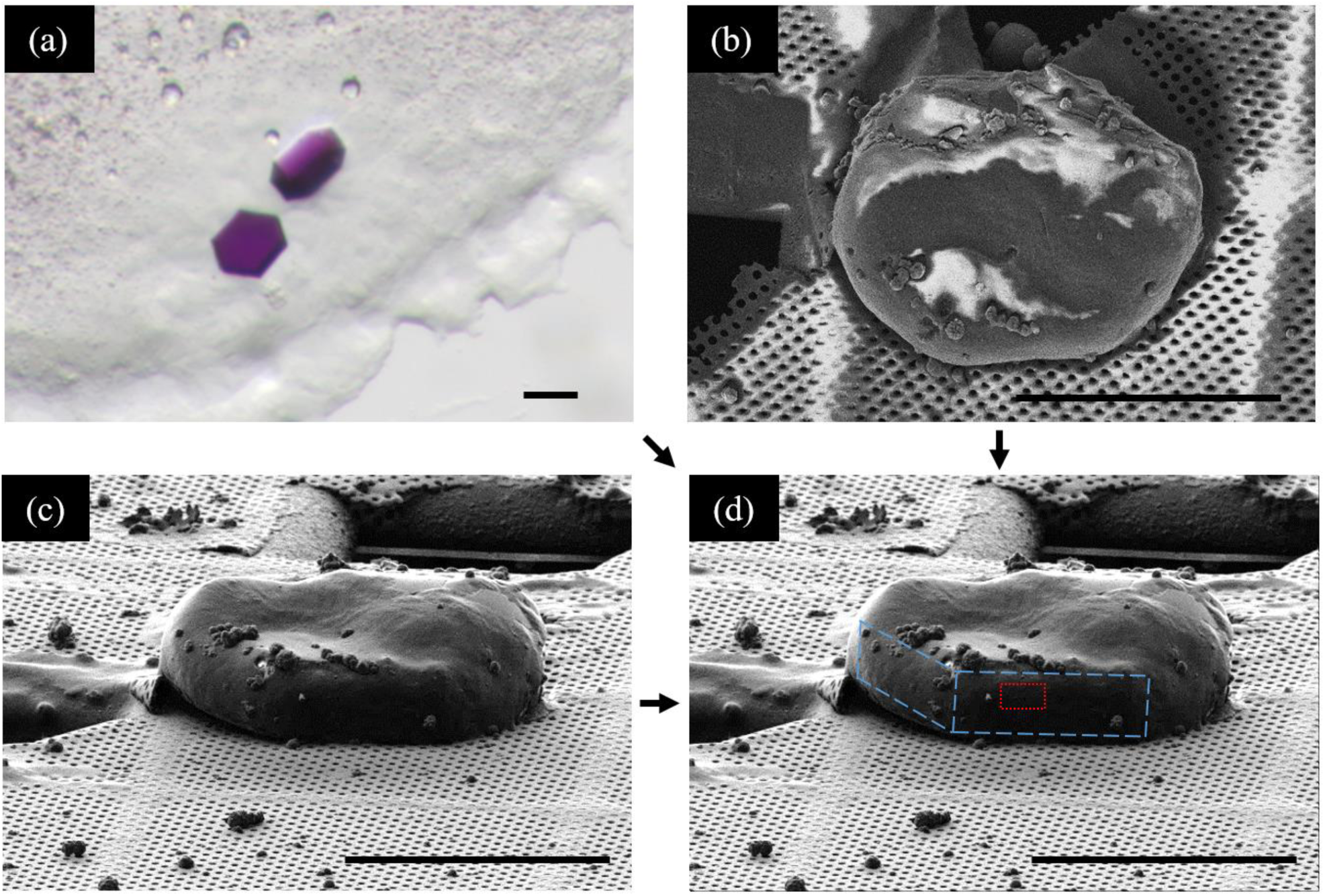
Images of bacteriorhodopsin crystals before cryo-FIB milling. Bright field optical micrograph of BR crystals grown in a LCP of monoolein at room temperature. SEM (b) and FIB (c) micrographs of a flash-frozen BR crystal with leftovers of the crystallization medium. (d) Geometrical features of the hexagon-shaped BR crystal are indicated by blue dash line on the FIB image. These features can be guessed by comparing (a), (b) and (c). The red dash line rectangle on (d) shows the selected area for further cryo-FIB milling. The scale bars correspond to 50 µm.

**Figure 3.**
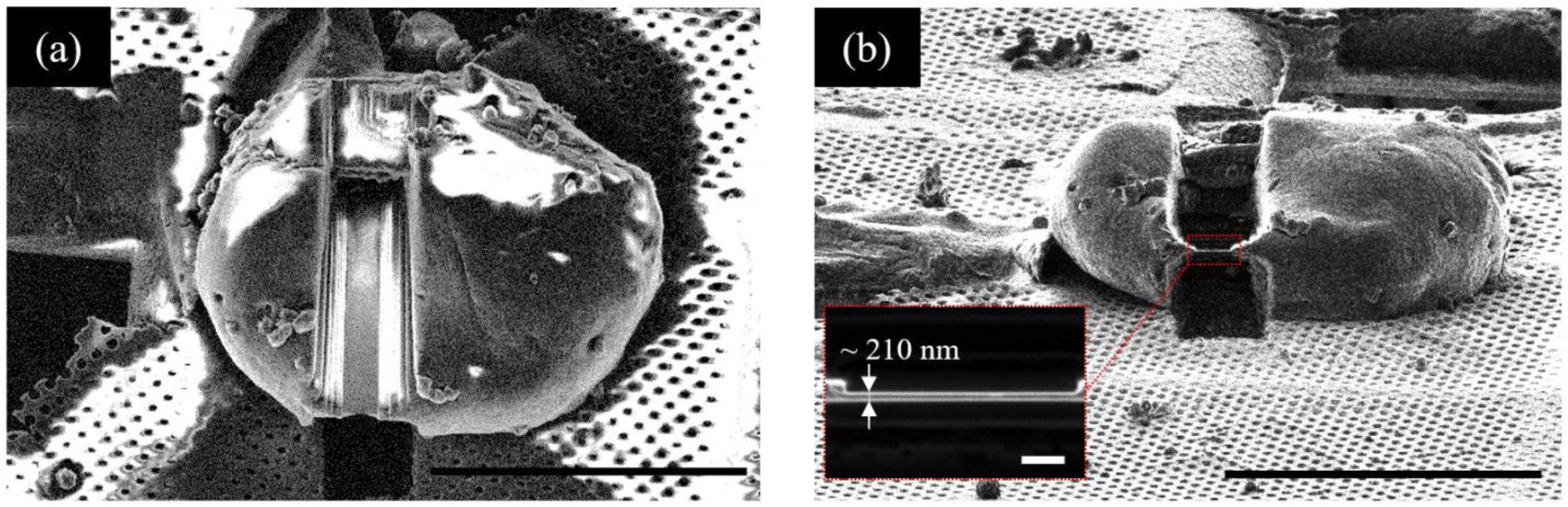
Images of the bacteriorhodopsin crystals after cryo-FIB milling. SEM image (a) and FIB image of the final lamella, obtained by the cryo-FIB milling. The inbox showing the zoomed image of the lamella. The scale bars correspond to 50 µm for (a), (b) and to 1 µm for the inset in (b).

### 2.3. Electron diffraction data collection

We used a Talos Arctica cryo-EM (Thermo Fisher Scientific), equipped with a Schottky field emission gun (XFEG) and operated at 200 kV corresponding to an electron wavelength of 0.02508 Å. The crystal lamella was identified in imaging mode (Fig. 4(a)) at a low magnification and at a low dose rate. The ED experiment was performed under parallel illumination conditions utilizing microprobe mode and spot size 11, resulting in an illuminated area of ∼4.6 µm in diameter on the sample plane. Under these conditions the flux for the electron beam was ∼0.006 e^−^/Å^2^/s (derived from a recorded fluence of ∼0.06 e^−^/Å^2^ during a 10-s exposure). ED data were collected from a zone of 1.4 µm in diameter as defined by a selected-area aperture. ED data were recorded as a still image with a CMOS-based Ceta 16M detector (Thermo Fisher Scientific), using 10-s exposure. The effective sample-to-detector distance was of 2767 mm, as calibrated using a standard gold-film sample.

**Figure 4.**
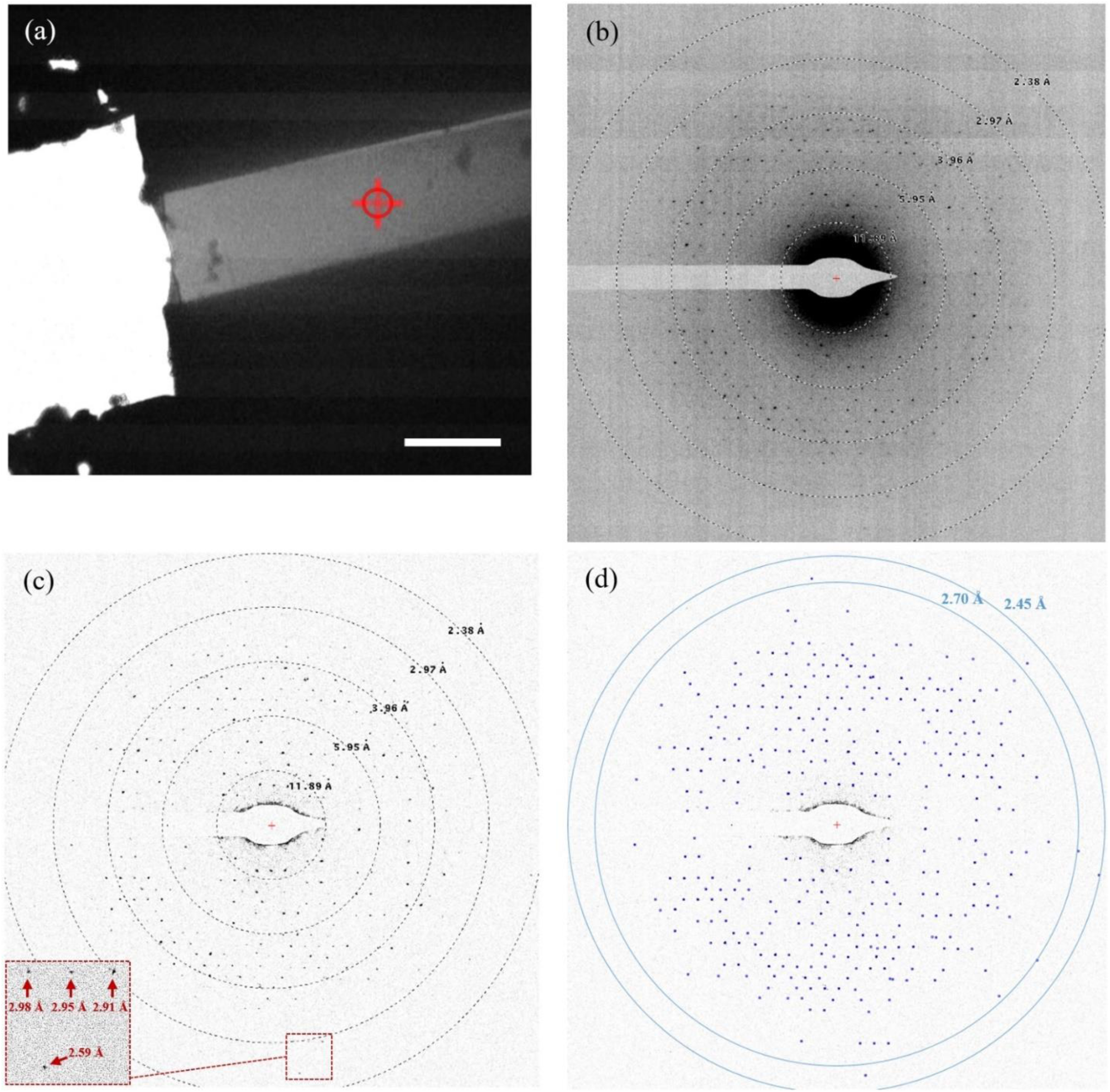
Electron diffraction experiment on a 210 nm thick lamella of bacteriorhodopsin, using a 200-kV cryo-TEM microscope. (a) TEM micrograph of the FIB-machined lamella of the BR crystal. Electron diffraction signal was collected from a 1.4-µm area of the lamella, indicated by a red circle with a cross. The scale bar on (a) corresponds to 5 µm. (b) The 200-kV electron diffraction pattern obtained from the area indicated in (a). (c) The electron diffraction image is corrected by subtraction of local moving-average background, calculated with the Adxv program (Adxv, 2013). The inset shows a close up of the electron diffraction pattern. (d) Diffraction peaks were automatically picked up by the Adxv software, and the diffraction peaks in the resolution shell of 2.7 Å - 2.45 Å had a mean signal-to-noise ratio <I/σ(I)> of 7.1.

## 3. Results and discussion

### 3.1. Crystal lamella preparation

Bacteriorhodopsin (BR) (Oesterhelt & Stoeckenius, 1971; Lanyi, 2004) was used as a model system, since it is one of the most well characterized MPs and had been instrumental for the development of a vast array of biophysical techniques, including *in meso* crystallization methods (Landau & Rosenbusch, 1996; Faham & Bowie, 2002; Takeda *et al*., 1998; Pebay-Peyroula *et al*., 1997), and used in cryo-electron microscopy and 2D electron crystallography of proteins (Henderson *et al*., 1990; Kimura *et al*., 1997; Mitsuoka *et al*., 1999; Subramaniam & Henderson, 2000). BR crystals were grown in a LCP of monoolein to a size of ∼60-80 µm in the longest dimension and to a thickness of ∼5-10 µm (Fig. 2(a)) by a standard LCP method (Pebay-Peyroula *et al*., 1997; Gordeliy *et al*., 2003; Polovinkin *et al*., 2014), which reproducibly produces the hexagon-shaped plate-like crystals of type I and space group P6_3_. One of the crystals was used for lamella preparation as shown schematically in Fig. 1.

Fig. 2 shows images of the crystals before milling. We were able to identify some of the edges of the hexagon-shaped BR crystal in the FIB image (Fig. 2(d)) despite the presence of leftovers of the frozen LCP matrix and the precipitant solution.

We expect that, in general, it would be a difficult task to find an appropriate area on the crystal to mill, if the navigation is based on FIB (or SEM) images, containing only the topographical information about the sample. Furthermore, locating crystals can be even more challenging, if all the geometrical features of the crystals are hidden in a bulky crystallization matrix. The problem likely becomes even more pronounced if the crystals are only a few microns in size or smaller, and are buried in tens of microns of crystallization matrix (e.g. see figures in (Liu *et al*., 2013, 2014; Zhu *et al*., 2019)). In principle, crystals can be localised in a bulky sample by correlative 3D light microscopies in a non-destructive way and with micron or submicron precision, prior to or in parallel to FIB milling. The localisation can be based on single- or multiphoton fluorescence (Judge *et al*., 2005; Svoboda & Yasuda, 2006; Madden *et al*., 2011; Leung & Chou, 2011; Jonkman & Brown, 2015; Li *et al*., 2015), second harmonic generation (SHG) (Kissick *et al*., 2010; Wampler *et al*., 2008) or Raman effect (Aksenov *et al*., 2002; Pezacki *et al*., 2011; Hazekamp *et al*., 2011; Opilik *et al*., 2013; Arzumanyan *et al*., 2016). Moreover, correlative cryo-fluorescence 3D light microscopy (cryo-FLM) is currently a part of the cellular cryo-ET workflow (Plitzko & Baumeister, 2019; Schaffer *et al*., 2019). This type of instrumentation and the methods outlined creates an efficient basis for site-specific cryo-FIB milling of bulky samples, containing target protein crystals.

Following the removal of excess crystallisation matrix, the selected part of the BR crystal (Fig. 3a) was thinned down to ∼210 nm thickness (Fig. 3c) using focused 30-kV Ga ions. The obtained thickness is within the optimal range, as recommended for ED signal optimizations (∼150-250 nm) in other 200-kV electron crystallography experiments with FIB-machined lysozyme crystal lamellae of various thicknesses (Zhou *et al*., 2019). It is worth noting that, considering the ED signal, 210 nm might not be the optimal thickness for BR crystals. In general, and also specific to the current case, the optimal thickness for ED data collection can be influenced by multiple factors, including: electron energy, which determines the mean elastic and inelastic paths and multiple scattering effects (Subramanian *et al*., 2015; Clabbers & Abrahams, 2018); effective thickness which is a function of tilt angle during data collection (Plitzko & Baumeister, 2019); mechanical stability for further handling and transportation (Plitzko & Baumeister, 2019; Zhou *et al*., 2019); the fact that a balance between the thickness-squared kinematic Bragg intensities and the thickness-dependent corruptions in intensities introduced by multiple scattering effects needs to be sought (Subramanian *et al*., 2015); and the bulk-solvent content of a protein crystal, which can influence the gross effect of multiple scattering (Michel, 1989; Weichenberger *et al*., 2015; Latychevskaia & Abrahams, 2019). Therefore, a thickness optimal for ED data accumulation can differ in each particular case. The easiest way of finding the optimal thickness could be to collect and to compare continuous-rotation ED data at different thicknesses for different electron energies, which is a goal set for future studies.

### 3.2. The electron diffraction experiment

Upon irradiating the 210-nm machined lamella (Fig. 3(b) and (c)) by a parallel 200-kV electron beam at a fluence of 0.06 e^−^/Å^2^ and accumulating the ED signal from the 1.4-µm diameter (Fig. 4a) zone, we observed an ED signal with an average signal-to-noise ratio <I/σ(I)> of 7.1 for peaks in the 2.7 Å - 2.45 Å resolution shell (Fig. 4). Considering the overall radiation dose limit of ∼2 e^−^/Å^2^ (for 200-kV electrons) as damage -”safe” (Hattne *et al*., 2018), we could roughly estimate that a wedge of around 15°-30° can be collected from a single BR crystal lamella, when continuously rotating it with a flux of 0.06 e^−^/Å^2^ per frame and a data collection speed of 0.5° - 1° per frame typically used for ED (Nannenga, 2020). Taking into account the fact that BR crystals belong to the space group P6_3_, one would expect that a ∼90°-wedge of reciprocal space needs to be collected to get a dataset resulting into a good completeness (Dauter, 1999; Borshchevskiy *et al*., 2011). This implies that, in order to collect ED data suitable for high-resolution structural determination with the experimental system used in the present article, at least 3-6 different lamellae have to be produced and have to survive transfer from the cryo-FIB instrument to the cryo-TEM microscope. Cryo-FIB milling is time consuming (∼5 h per lamella in our case) and the transfer-surviving rate was about ∼50% in our case (see also (Zhou *et al*., 2019; Plitzko & Baumeister, 2019)), making the overall sample preparation cumbersome.

We note that the experimental system that was available to us was not optimized for diffraction experiments. In particular, the CMOS-based detector Ceta 16M (Thermo Fisher Scientific) had a quantum efficiency of only 0.09 at 1/2 Nyquist frequency. Direct electron detectors have much higher sensitivity (McMullan *et al*., 2016; Naydenova *et al*., 2019; van Genderen *et al*., 2016) and would allow significant improvements in data quality and resolution, allowing experimenters to collect data over a much wider angular range and at much improved signal to noise ratios (Hattne *et al*., 2019). Such detectors would permit higher resolution ED data while also minimizing the number of lamellae necessary to reach good completeness.

In another experiment, using a BR crystal lamella similar to the one reported here but of a lower quality (diffraction to about 4 Å), we tried to collect ED diffraction data over a 20°-wedge of reciprocal space in continuous rotation mode. The unit cell dimensions were a = b = 61.2 Å, c = 109.2 Å; α = β = 90°, γ = 120°, when fixing the space group P6_3_ and using the data reduction software XDS (Kabsch, 2010). This result is consistent with values observed in X-ray diffraction experiments (Luecke *et al*., 1999; Borshchevskiy *et al*., 2011; Polovinkin *et al*., 2014).

### 3.3. Practical aspects of lamella preparation

We found that it was important to use a low FIB current of 10 pA at the final step of milling when removing the last ∼100 nm from each side of the lamella (see section 2.2). In a few experiments, when we tried to minimize the FIB-milling time, we used a 30-pA current instead of 10-pA one in the final step. Although the milling time decreased from ∼60 min to ∼20 min, the use of the higher current led to lower quality lamellae, diffracting to 7 - 4 Å only. Of course, this could have been caused by other factors such as crystals grown to lower quality or damaged during transfer from crystallization probe to a TEM grid, nevertheless, it is worth mentioning that similar observations were reported by (Martynowycz *et al*., 2019*b*) in experiments on FIB-milled lamellae of proteinase K crystals. The authors utilized a cryo-FIB instrument similar to the one used in our present study and showed that focused Ga^+^ beams with 10-pA and 30-pA current produced lamellae of different qualities, diffracting to 1.8 Å and 2.1 Å, respectively. We can speculate that local heating induced by higher currents can provoke disordering or devitrification processes, which are less pronounced in the case of lower currents (see (Marko *et al*., 2006; Plitzko & Baumeister, 2019)). These and similar observations suggest that milling conditions at cryogenic temperatures should be mapped out for each type of crystal to find a suitable balance between diffraction quality and overall milling time. Crystals of macromolecules can have significantly different water content and crystallisation medium (Michel, 1989; Weichenberger *et al*., 2015), and this may influence the results. FIB-milling can be significantly accelerated by using other ions than Ga^+^. For example, new plasma FIBs utilizing noble gas ions promise removal rates that are up to 60x faster than the rate achievable with Ga ions (Marko *et al*., 2006). The potential of utilizing FIBs with different ions for macromolecular electron crystallography has yet to be explored.

## 4. Conclusion

Sample preparation is a critical and onerous step in both electron microscopy and crystallography of macromolecules. We have presented a sample preparation scheme for electron crystallography for studies on 3D crystals of membrane proteins grown *in meso*, and using FIB milling under cryogenic conditions. Our results demonstrate that a lamella of high quality for ED studies can be prepared from a 3D MP crystal surrounded by a viscous and sticky lipidic mesophase. The cryo-FIB milling technique gives a precise control over the thickness of crystal lamellae, which potentially permits ED data optimization in terms of resolution and influence of multiple scattering. This can lead to more accurate kinematic structure factors and the precise location of ions and hydrogen atoms in the electrostatic potential maps (Yonekura *et al*., 2015; Clabbers *et al*., 2019), that can be crucial for understanding vital processes run by MPs.

We believe that the cryo-FIB milling method can be applicable to crystals of membrane or non-membrane proteins grown or embedded in various lipidic mesophases, including bicells and vesicles, or in other viscous crystallization media such as concentrated polyethylene glycol (PEG) solutions and media used for high-viscosity injectors (e.g. media used in (Wolff *et al*., 2020)). Moreover, we believe that, the cryo-FIB milling technique can also be suitable for crystals of few microns or even of smaller sizes. 3D electron micro-/nano-crystallography of MP crystals grown *in meso* can be facilitated by correlative 3D light microscopies, which can help to localize very small crystals in a bulky crystallization environment for subsequent cryo-FIB milling.

## Acknowledgements

We thank V. Gordeliy (Institut de Biologie Structurale (IBS), France; Forschungszentrum Jülich GmbH, Germany; Moscow Institute of Physics and Technology, Russia) for kindly providing bacteriorhodopsin-containing purple membranes from *Halobacterium salinarum* S9 cells. We are grateful to Jiri Pavlicek and Jan Stransky (BIOCEV CF Crystallization of Proteins and Nucleic Acids) and Jiri Novacek and Jana Moravcova (CEITEC CF Cryo-electron Microscopy and Tomography) for experimental support.

